# Evolutionary analysis of Quinone Reductases 1 and 2 suggest that NQO2 evolved to function as a pseudoenzyme

**DOI:** 10.1101/2022.11.11.516182

**Authors:** Faiza Islam, Nicoletta Basilone, Eric Ball, Brian Shilton

## Abstract

Quinone reductases 1 and 2 (NQO1 and NQO2) are paralogous FAD-linked enzymes found in all amniotes. NQO1 and NQO2 have similar structures and can both catalyze reduction of quinones and other electrophiles. The two enzymes differ in their cosubstrate specificity, with NQO1 using cellular redox couples NAD(H) and NADP(H), while NQO2 is almost completely inactive with these cosubstrates, and instead uses dihydronicotinamide riboside (NRH) and small synthetic cosubstrates such as *N-*benzyl-dihydronicotinamide (BNAH). We used ancestral sequence reconstruction to investigate the catalytic properties of a predicted common ancestor and 2 additional ancestors from each of the evolutionary pathways to extant NQO1 and NQO2. In all cases, the small nicotinamide cosubstrates NRH and BNAH were good cosubstrates for the common ancestor and the enzymes along the NQO1 and NQO2 lineages. In the case of NADH, however, extant NQO1 evolved to a catalytic efficiency 100x higher than the common ancestor, while NQO2 has evolved to a catalytic efficiency 1000x lower than the common ancestor. In addition, 13 chimeric enzymes were created to investigate the molecular basis of cosubstrate specificity, which was further elaborated by site-directed mutagenesis of the ancestral NQO2. Overall, the results suggest a selective pressure for evolution of NQO1 towards greater efficiency with NADH, and for NQO2 towards extremely low efficiency with NADH. These divergent trajectories have implications for the cellular functions of both enzymes, but particularly for NQO2 whose cellular functions are only beginning to be uncovered.

## Introduction

Quinone reductases comprise an ancient family of flavoenzymes that afford the two-electron reduction of quinones and other electrophiles as a detoxification strategy. These enzymes use a catalytic FAD or FMN to receive a hydride from reducing cosubstrates NAD(P)H and then transfer the hydride to the quinone substrate or another electrophile (Deller, Macheroux, and Sollner 2008; Vasiliou, Ross, and Nebert 2006). Humans have two paralogous quinone reductase genes: Quinone Reductase 1, *nqo1* and Quinone Reductase 2, *nqo2* (Jaiswal et al. 1990; Zhao et al. 1997). The two proteins share 47% sequence identity and a strikingly similar overall structure. NQO1 has numerous roles, and its regulation as part of the Keap1-Nrf2 pathway provide a clear enzymatic function in the cellular response to oxidative stress (Lind, Hochstein, and Ernster 1982; Sihvola and Levonen 2017; Ross and Siegel 2021).

The properties, regulation, and potential functions of NQO2 have been recently reviewed (Janda et al. 2020). While the cellular functions of NQO2 are still somewhat of a mystery (S Liao, Dulaney, and Williams-Ashman 1962; Vella et al. 2005), there has been intriguing progress in recent years. In the brain, NQO2 was found to be overexpressed in a model system for age-related memory impairment and a pharmacological model of amnesia (Brouillette et al. 2007), and NQO2-knockout mice demonstrate enhanced learning (Benoit et al. 2010). Roles for NQO2 in learning and memory, along with insight into the cellular mechanisms, have been investigated by Rosenblum and co-workers (Rappaport et al. 2015; Gould, Sharma, et al. 2020; Gould et al. 2021), with new evidence that NQO2 dysfunction may contribute to metabolic stress and neurodegeneration (Gould et al. 2023). Our own interest in NQO2 began with the observation that it was an off-target interactor with inhibitors of the kinase CK2 (Duncan et al. 2008; Kevin K. K. Leung and Shilton 2015). At the time, NQO2 was known to be an off-target interactor with two other kinase-targeted therapeutics, imatinib, targeted to the Abl kinase, and bisindolylmaleimide inhibitors targeted towards PKC (Bantscheff et al. 2007; Rix et al. 2007; Winger et al. 2009; Brehmer et al. 2004). The broad extent of NQO2 off-target interactions with kinase inhibitors was demonstrated in a comprehensive screen of over 200 clinically used kinase inhibitors, where NQO2 was found as an off-target interactor with over 30, by far the most frequent non-kinase interactor discovered (Klaeger et al. 2017). This finding suggests that NQO2 may play a role in the cellular effects of these drugs, such that interaction with NQO2 is selected for during drug screening and optimization. It is notable that unlike NQO2, NQO1 has not been picked up in off-target screens for kinase-targeted drugs. NQO2 has also been found to interact with antimalarials (Graves et al. 2002; Kwiek, Haystead, and Rudolph 2004). Until the cellular functions of NQO2 are understood in greater detail, it is difficult to know how its interactions with these various drugs might affect cell metabolism, signalling, and viability.

NQO2 is unusual because it cannot efficiently use the cellular redox couples NAD(H) and NADP(H) (Shutsung Liao and Williams-Ashman 1961; S Liao, Dulaney, and Williams-Ashman 1962; Chen, Wu, and Knox 2000; Islam et al. 2022). Intriguingly, NQO2 can efficiently use various smaller dihydronicotinamide cosubstrates such as dihydronicotinamide riboside (NRH) and *N-*propyl-dihydronicotinamide (Shutsung Liao and Williams-Ashman 1961; S Liao, Dulaney, and Williams-Ashman 1962), *N-*benzyl-dihydronicotinamide (BNAH), *N-*methyl-dihydronicotinamide (NMEH) and others (Knox et al. 2000) as sources of electrons. Of these smaller reducing cosubstrates, only NRH, and its oxidized form, nicotinamide riboside (NR) function in cells where they contribute to salvage pathways to regenerate NAD^+^ (Bieganowski and Brenner 2004; Yang et al. 2019; 2020; Giroud-Gerbetant et al. 2019). However, there is currently no evidence that the relatively low levels of NR and NRH can support enzymatic activity of NQO2 by functioning as a cellular redox couple like NAD(H) and NADP(H). In fact, experiments in cells using a substrate, CB1954, that becomes highly toxic upon NQO2-catalyzed reduction, indicate that NQO2 can only function effectively as a redox enzyme when it is supplied with exogenous small nicotinamide cofactors such as NRH or BNAH (Knox et al. 2000; Islam et al. 2022).

Why would an enzyme evolve such an unconventional cosubstrate specificity? The cosubstrate specificity of human NQO2 is conserved in other mammals (S Liao, Dulaney, and Williams-Ashman 1962; Zhao et al. 1997; Celli et al. 2006). More recently, the inability of NQO2 to efficiently use NAD(P)H was demonstrated for reptile (*Alligator mississipiensis*) and bird (*Anas platyrhynchos*) NQO2, indicating that this unusual property of NQO2 is conserved throughout the amniotes (Islam et al. 2022). A phylogenetic analysis of NQO1 and NQO2 sequences pointed to a gene duplication event in a vertebrate ancestor before the split of the ray-finned and lobe-finned fish, approximately 460 million years ago (MYA) (Islam et al. 2022; Hedges 2009). Gene duplication is a significant force for functional diversity in protein families (Ohno 1970; Zhang 2003). The most common fate of duplicated genes is the degeneration and silencing of one of the pair to a non-functioning pseudogene (Lynch and Conery 2000; McGrath et al. 2014). In less common cases, both copies of the gene could be maintained if they confer a survival benefit (Walsh 2003). In these cases, there are two possible fates for the duplicated genes: they could develop sub-functionalizations, where the original function is partitioned across both copies; alternatively there could be neo-functionalization, where one copy acquires a new function (Zhang 2003). The difference in cosubstrate specificity observed between NQO1 and NQO2 could indicate either sub-or neo-functionalization. In this regard, an important question is whether NQO2 evolved to use NRH efficiently, or whether it evolved specifically *not* to use NAD(P)H.

We have used ancestral sequence reconstruction to investigate the evolution of cosubstrate specificity in NQO1 and NQO2. Ancestral sequence reconstruction has been used to trace the molecular mechanisms of substrate or inhibitor specificity for many enzyme families (Merkl and Sterner 2016; Wilson et al. 2015; Hobbs et al. 2012; Harris et al. 2022). We have reconstructed the predicted common ancestor of NQO1 and NQO2, along with two ancestors of each enzyme along their evolutionary paths from the common ancestor. The reconstructed NQO enzymes are catalytically active with the common quinone substrate, menadione. However, kinetic analyses show that after duplication from the common ancestor, NQO2 lost the ability to efficiently use NADH as a cosubstrate, whereas the catalytic efficiency of NQO1 improved compared to the common ancestor. In contrast to the consistent changes towards increased (NQO1) or decreased (NQO2) catalytic efficiency with NADH, all of the enzymes were able to use BNAH or NRH with moderate to high efficiency. This investigation into the evolution of the two enzymes indicates that NQO2 has evolved to avoid efficient reduction by NADH and NADPH rather than use NRH efficiently, and has provided insight into the specific molecular determinants of the difference in cosubstrate specificity between NQO1 and NQO2.

## Materials and Methods

### Reagents

Ampicillin, FAD, NADH, and NADPH were purchased from Sigma-Aldrich. 1-benzyl-1,4-dihydro-3-pyridinecarboxamide (BNAH) was purchased from TCI Chemicals. Nicotinamide riboside (NR) chloride was purchased from Chromadex (Los Angeles, CA) as “Tru Niagen” nutritional supplements. To reduce the NR to NRH (dihydronicotinamide riboside), the contents of each capsule (263 mg of nicotinamide riboside) were dissolved in 10 mL of 1 M NaHCO_3_, pH 8.1, and the solution was filtered to remove insoluble microcrystalline cellulose. Following the basic protocol outlined in Zarei *et al*. (Zarei et al. 2021) dihydronicotinamide riboside (NRH) was prepared by degassing the NR solution, adding 2.7 equivalents of Na_2_S_2_O_4_ and incubating under nitrogen for 3 hours. NRH was purified by reversed-phase HPLC using a semi-preparative Zorbax XD8-C18 column (9.4 x 250 mm; Agilent).

### Evolutionary Analysis

The evolutionary analysis consisted of multiple sequence alignments of NQO1 and NQO2 sequences that were found using human NQO1 and NQO2 protein sequences as bait against the database of non-redundant protein sequences using the NCBI BLASTp server. Whether a given sequence is NQO1 or NQO2 was determined by the annotation suggested by NCBI, by the percentage identity, and by the presence or absence of the 43-residue C-terminal domain of NQO1. A total of 172 sequences were collected. A preliminary multiple sequence alignment was calculated with MUSCLE (Edgar 2004) and showed that the sequences could be aligned with minimal gaps or insertions. The next stage of the evolutionary analysis was ascertaining the evolutionary relationship between the protein sequences by constructing a phylogenetic tree. Simultaneous multiple sequence alignment and phylogenetic tree construction were done using Phylobot (http://www.phylobot.com; Hanson-Smith and Johnson 2016).

The phylogenetic tree was constructed by maximum likelihood with the JTT+CAT evolutionary model (Jones et al 1992). Finally, the multiple sequence alignment and the phylogenetic tree were used to infer the amino acids of the sequences at the nodes of the phylogenetic tree. The predicted protein sequences from Phylobot at specific nodes of interest were selected for analysis. The resurrected sequences comprised the amino acid residues with the highest probability at each position. The amino acid sequences were reverse-translated to yield the corresponding DNA sequence.

### Cloning of constructs and protein expression

Cloning of constructs was carried out using *E. coli* XL-1 Blue competent cells. Ancestral DNA sequences were ordered as genes from Integrated DNA Technologies (“IDT”, Coralville, Iowa). The sequences were codon-optimized for expression in bacteria using IDT-provided tools. Two restriction sites were engineered into the sequences, KasI/Ehe1 at the 5’ end, and HindIII at the 3’-end to facilitate cloning into the pProEx-HTA expression plasmid (Invitrogen) in a way that fuses a coding sequence for a 6xHis tag followed by a TEV cleavage site (i.e., HHHHHHDYDIPTTENLYFQ/GA) to the 5’ end of the quinone reductase gene. Proteolytic cleavage with TEV protease removes the affinity tag, leaving only an extra GA sequence at the N-terminus of the proteins.

The chimeric sequences were constructed from CA2NQO2 and CA2NQO1 by Gibson Assembly according to previously published protocol (Ball and Basilone 2022). Overlapping PCR products encoding fragments of the CA2NQO1 and CA2NQO2 were assembled by Gibson assembly and inserted into a modified pET vector, pEBC, via Gibson Assembly. The modified pEBC plasmid contains the coding sequence for fusion of a cellulose-binding module (CBM) coding sequence to the 5’ end of the inserted gene, to produce a fusion construct with a CBM at the N-terminus for small-scale affinity purification, separated from the inserted gene product by a Tobacco Etch Virus (TEV) protease cleavable linker. BamHI and XhoI restriction sites were used to open the plasmid vector. Primers for the target sequences were designed to incorporate the cut sites and some upstream sequences matching the vector to provide for overlapping ends during Gibson Assembly.

Cloning of the NQO2 mutants was also done with the same Gibson Assembly strategy as the chimeras. PCR primers were ordered to either make a short NQO2 mutant without the C-terminal tail or a mutant with the longer 43-residue C-terminal of CA2NQO1. The sequence of all constructs was verified with Sanger sequencing at the Robarts Research Institute, University of Western Ontario.

### Protein Expression and Purification

Protein expression was done in *Escherichia coli* BL21-DE3 strains under ampicillin selection. For large-scale protein production, 4 1L cultures were inoculated with a single colony starter culture and grown overnight at 37 °C in Studier’s ZYP-5052 auto-induction media (Studier 2005). The ancestral proteins were purified by previously described methods (Kevin Ka Ki Leung, Litchfield, and Shilton 2012; Islam et al. 2022). Bacterial cell pellets were thawed, treated with DNase and lysozyme, and dispersed by the Dounce homogenizer and then the French press. The cell lysates were supplemented with 50 mM Na-phosphate, pH 7.4 and 1 M NaCl and 25 mM imidazole. The cells were ultracentrifuged at 4°C at 100,000 x g for 1 hour and 20 minutes to remove cell membranes and debris. After centrifugation, the pellet was discarded. The supernatant was applied to a 5 mL Ni^2+^-NTA affinity (IMAC Sepharose, Cytiva) equilibrated with the same buffer. The 6xHis-tagged protein was eluted with 250 mM imidazole in 50 mM Na-phosphate, 1 M NaCl, pH 7.4. After Ni^2+^-affinity purification, the protein was dialyzed against 100 mM Tris-HCl, 0.5 M NaCl, 5 mM DTT, and 5 mM EDTA, pH 8.2. The dialyzed and partially purified protein was then digested with TEV protease to remove the affinity tag.

For anion-exchange chromatography of acidic proteins related to NQO2, the proteins were dialyzed against 10 mM Tris, 1 mM EDTA, 0.5 mM TCEP (tris(2-carboxyethyl)phosphine), pH 8.4 in preparation for anion exchange chromatography. The protein was applied to a 1.6 x 15 cm column of Q-Sepharose HP (Cytiva) equilibrated with 20 mM Tris, 1 mM EDTA, and 1 mM TCEP at pH 8.4. The column was developed with a gradient from 0 to 500 mM NaCl at a flow rate of 2 mL/minute while collecting 8 mL fractions. For cation-exchange chromatography of basic proteins more closely related to NQO1, the protein was dialyzed against 20 mM Na-phosphate, 0.5 mM TCEP, and 1 mM EDTA, pH 5.0. The dialyzed protein was applied to a 1.6 x 15 cm column of S-Sepharose HP (Cytiva) equilibrated with 20 mM NaP, 1 mM EDTA, and 0.5 mM TCEP, pH 5.0. The column was developed with a gradient of from 0 to 500 mM NaCl at flow rate of 2 mL/min flow while collecting 8 mL fractions. After ion-exchange chromatography, the protein was pooled and concentrated by ultrafiltration (Amicon) for gel-filtration chromatography. For FAD reconstitution, the concentrated protein samples were incubated with 1 to 3 M Guanidine-HCl, 10 mM FAD, 1 mM ZnCl, and 10 mM DTT at 4°C. After gentle mixing, the protein samples were centrifuged to remove any precipitate and then applied to a 2.6 x 65-cm column of Superdex-200 preparatory grade gel filtration resin (Cytiva), equilibrated with 50 mM Tris, pH 8.0 and 150 mM NaCl, 10 µM FAD, 10 µM Zn^2+^. After gel filtration, NQO2 was dialyzed against 50 mM HEPES, 100 mM NaCl, pH 7.4 and concentrated by ultrafiltration using an Amicon stirred cell. The purified and concentrated proteins were flash-frozen in liquid nitrogen and stored at −80 °C. Protein concentration was determined with the Bradford assay. To assess FAD content, we measured the UV/Vis absorbance spectra of the proteins at the concentration (0.3 – 1.5 mg/mL) in 10 mM HEPES, pH 7.4, from a wavelength of 600 nm to 250 nm with the Cary 100 spectrophotometer.

Small-scale protein expression of the chimeric CBM-fusion constructs was done according to previously described protocols (Ball and Basilone 2022). For functional screening of the chimeric constructs, BL21(DE3) cells were transformed with the expression vector for the CBM-fusion, carrying an ampicillin marker, and a pLysS plasmid that constitutively expressed lysozyme and carries a chloramphenicol marker. One or two transformed colonies were picked and grown overnight at 37°C in 2.5 mL of Terrific Broth that included ampicillin (100 µg/mL), chloramphenicol (34 µg/mL) and IPTG (0.2 mM). Cells were harvested from 1.5 mL of the cultures and resuspended in 0.75 mL of 10 mM Tris-HCl, pH 7.5. The cells were lysed by two freeze-thaw cycles followed by a brief five-second sonication. The lysed samples were centrifuged at 20,000 x g for 15 minutes to remove cell debris. The resulting supernatant (cell extract) was added to one well of a six-well tissue-culture plate and supplemented with 10 mM Tris-HCl, 1 mM EDTA, pH 7.5 to bring the total volume to 2 mL. For small-scale affinity purification, Whatman #1 filter paper was used to make 6mm diameter paper disks using a hole-punch; 4 disks were added to each of the cell extracts and the 6-well plate incubated overnight at 4℃ to allow adsorption of the CBM-fusion protein. The disks were washed with TEN100 buffer (10 mM Tris-HCl, 100 mM NaCl and 1 mM EDTA, pH 7.5) using a sintered glass funnel under vacuum. To elute the protein off the disks, they were transferred to a 200 µL microcentrifuge tube with a small hole in the bottom, which in turn was placed inside a 500 µL microcentrifuge tube. This assembly was centrifuged to remove excess liquid and 28 µL of TEV protease solution (100 μg/mL TEV protease in TEN100 buffer containing 5 mM DTT) was added to the disks, which incubated at room temperature for 1 hour. The liberated protein was collected from the disk by centrifugation. The disks were washed with an additional 28 µL TEN100. The protein concentration was determined from an aliquot by TCA precipitation and Lowry protein assay.

### Thermal Shift Assay

To examine the stability of the constructs, the melting temperature (T_m_) of the proteins was determined with the Applied Biosystems Real-Time PCR System using the QuantStudio^TM^ 3 Real-Time PCR System. The thermal shift assay works by monitoring the change in fluorescence of the Sypro Orange Dye^TM^, with an excitation maximum of 472 nm and an emission maximum of 570 nm. The PCR system thermally denatures proteins by changing the temperature in a controlled manner. The fluorescence emission from Sypro Orange^TM^ dye increases when it binds to the exposed hydrophobic regions of unfolded proteins. Proteins, 0.2 to 0.3 mg/mL in 100 mM HEPES, pH 7.4 and 200 mM NaCl, 5 mM DTT were incubated with 1x Sypro Orange Dye^TM^. Each protein was investigated in at least four replicates in a total volume of 20 μL in a 96-well plate. As negative controls, the fluorescence of dye only and buffer only were measured. As FAD excitation and emission overlap with Sypro Orange^TM^, the change in fluorescence of FAD from the protein alone was measured as a control. There was no comparable signal from the FAD alone at the protein concentrations used, so it was likely not to affect the rest of the assay.

The 96-well plate was incubated on ice before starting the thermal denaturation assay. To generate the melt curves, the temperature was increased to 25°C at a rate of 1.6 °C/s and then held at 25°C for 2 minutes. Then, fluorescence data collection was started, and the temperature was increased to 99.0 °C at a rate of 0.05 °C/s. The temperature was then held at 99.0 °C for 2 minutes, after which the assay was finished. The Protein Thermal Shift^TM^ Software v1.0 analyzed the melt curve data.

### Steady-State Kinetics

We used a previously described protocol to determine the enzymatic activity of all five ancestral proteins (S Liao, Dulaney, and Williams-Ashman 1962). For kinetic assays with BNAH and menadione, the stock BNAH and menadione solutions in methanol were prepared fresh every day and kept on ice during the duration of the kinetic experiments. The concentration of each reagent was determined spectroscopically using the extinction coefficient ε_355 nm_ of 7220 M^-1^ • cm^-1^ for BNAH and ε_333 nm_ of 2450 M^-1^ • cm^-1^ for menadione. The kinetic reactions were conducted by monitoring the decrease in absorbance at 355 nm using a Cary 100 spectrophotometer to measure the rate of oxidation of the reducing cosubstrate. Enzyme assays were carried out for 2 minutes in 1 mL volumes in 100 mM HEPES, pH 7.4 at 25 °C under constant stirring with a small magnetic stir bar. Enzymatic activity was measured for a range of BNAH (500 µM - 5 µM) and menadione (100 µM - 2 µM) concentrations, using an appropriate protein concentration from 0.5 nM to 0.5 µM. Each reaction was carried out in triplicate or greater.

For the kinetic assays with NADH, the same protocol was used. Stock NADH was dissolved in 100 mM Tris-HCl, pH 8.0, and prepared fresh before use. NADH (10 µM to 2 mM) oxidation was measured by monitoring the change in absorbance at a wavelength of either 360 nm for high NADH concentrations (ε_360 nm_ of 4147 M^-1^ • cm^-1^) or at 340 nm (ε_340 nm_ of 6220 M^-1^ • cm^-1^) for lower concentrations of NADH. For the kinetic assays with NRH, the cosubstrate was dissolved in 100 mM Tris-HCl, pH 8.0 and concentrations from 10 µM to 2 mM were used in the assays. The reaction progress was monitored by measuring the change in absorbance at a wavelength of 360 nm (ε_360 nm_ of 4360 M^-1^ • cm^-1^) or at a wavelength of 380 nm (ε_380 nm_ of 1070 M^-1^ • cm^-1^). The initial rates were globally fit to a ping-pong bi-substrate Michaelis Menten equation (Equation 1, below) to obtain *K*_M_ values for both substrates and *K*_I_ for menadione using Prism (www.graphpad.com).

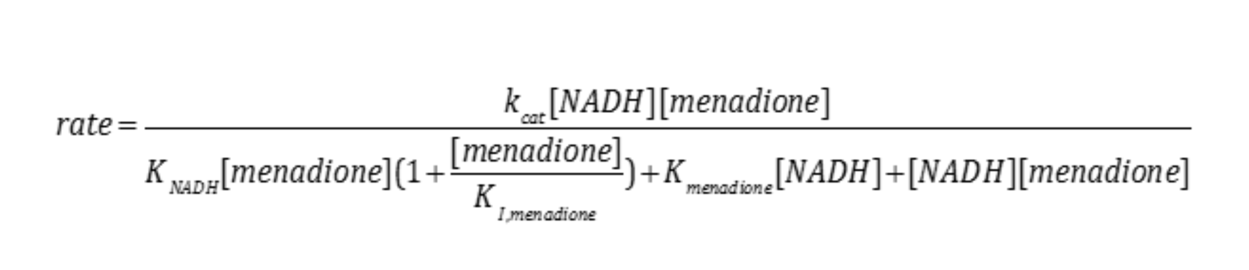

## Results

### Conservation of NQO2 co-substrate specificity among amniotes

An initial evolutionary analysis of NQO1 and NQO2 showed that NQO1 and NQO2 orthologs were present in all tetrapod genomes, except for some amphibians, and that the unusual cosubstrate specificity of NQO2 was conserved in a reptile and bird, and therefore likely conserved among all amniotes (Islam et al. 2022). This observation raises two possibilities for natural selection and evolution of NQO2. The first is that efficient use of NRH was subject to natural selection, and during this process the ability of NQO2 to use NAD(P)H was lost. A second alternative is that NQO2 has evolved to be unable to use NAD(P)H efficiently; that is, NQO2 may have gained a non-enzymatic cellular function that is incompatible with an ability to efficiently accept electrons from NAD(P)H. Deciding between these two alternatives will help to elucidate the cellular functions of NQO2.

### Resurrection of ancestral NQO1 and NQO2 proteins

A phylogeny based on extant NQO1 and NQO2 sequences indicated a gene duplication event in an ancestral fish approximately 450 MYA, prior to the divergence between the ray-finned and lobe-finned fishes; an abbreviated phylogenetic tree is illustrated in Figure 1A with the full phylogenetic tree in the Supplemental Figure S1. To track the evolution of NQO1 and NQO2, we reconstructed their common ancestor and two additional enzymes along each of the evolutionary pathways to extant NQO1 and NQO2. The ancestral sequence reconstruction method relies on multiple sequence alignment of extant related protein sequences and their phylogenetic relationship to reconstruct the ancestral protein sequences at the nodes of the phylogenetic tree. Ancestral NQO1 and NQO2 sequences were predicted using 172 extant NQO1 and NQO2 sequences and a combined optimization of the sequence alignment and phylogeny by maximum likelihood using PhyloBot (Hanson-Smith and Johnson 2016; Figure 1 and S1). This phylogenetic tree closely matched the accepted species-level tree of life (Hedges 2009). The phylogenetic relationship of NQO sequences is consistent with a previous analysis of quinone reductase sequences, with NQO1 and NQO2 arising from a gene duplication and forming separate branches of the tree (Vasiliou, Ross, and Nebert 2006).

**Figure 1.**
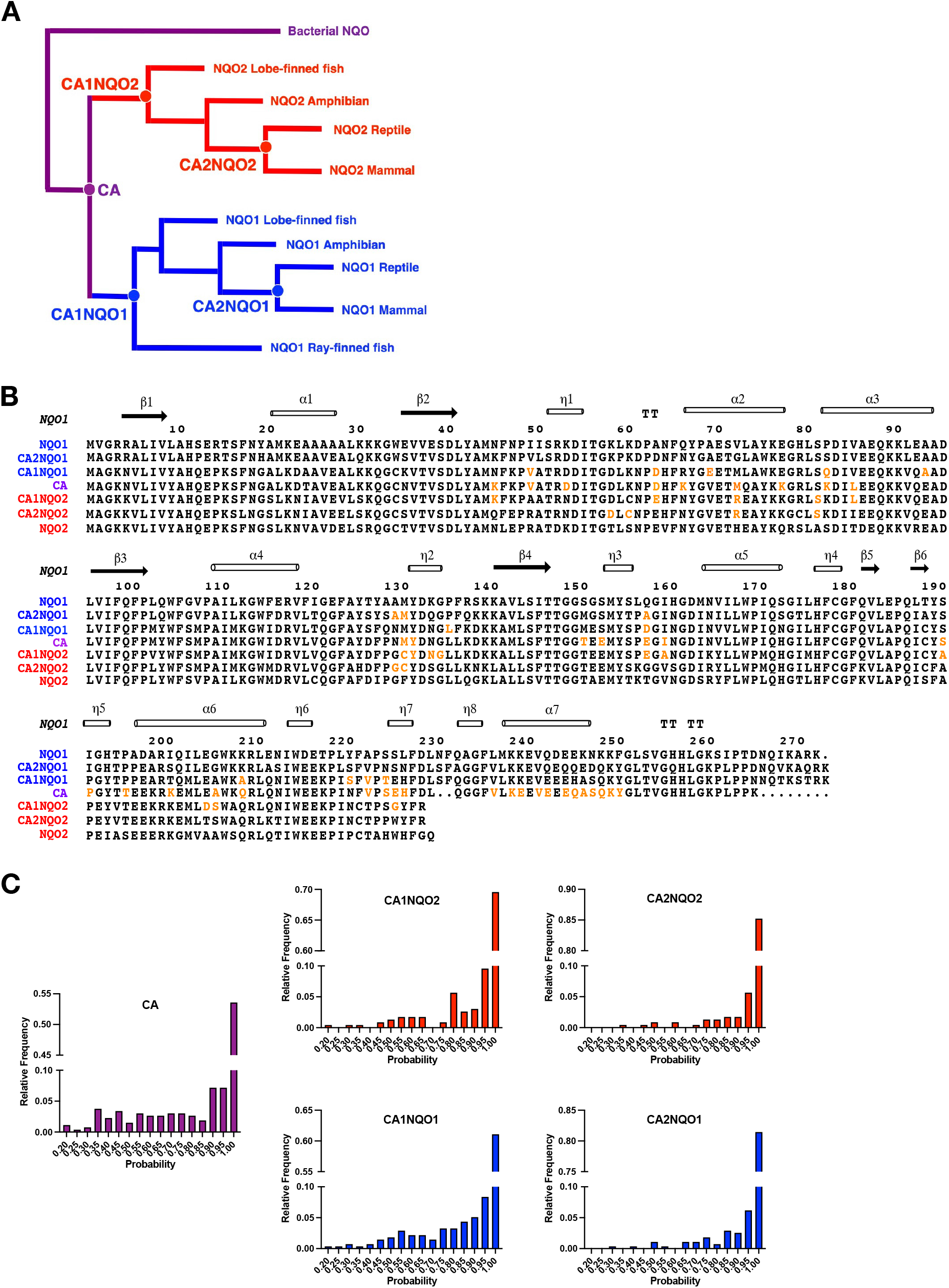
Prediction of Ancestral Quinone Reductase Sequences. (A) Summary of the phylogenetic tree for NQO1 and NQO2; node points indicated with circles are sequences that are reconstructed in this study. CA or “common ancestor” was the ancestral gene that was duplicated to yield extant NQO1 and NQO2. Node points CA1NQO1 and CA1NQO2 represent the predicted sequences at the top of the NQO1 and NQO2 phylogenetic trees. Node points CA2NQO1 and CA2NQO2 are at the top of the NQO1 and NQO2 phylogenetic trees for the amniotes. The complete phylogenetic tree is provided in Supplemental Figure S1. (B) Multiple sequence alignment of NQO1 and NQO2 along with the maximum likelihood sequences of the resurrected ancestral proteins. The residues in orange were predicted with probability below 0.6. The secondary structure for extant human NQO1 is provided at the top of the alignment, with α, β, and η representing alpha-helices, beta strands, and 3_10_ helices, respectively; “TT” indicates β-turns. NQO1 and NQO2 have similar overall structures, and therefore the secondary structure for NQO2 follows closely that of NQO1. (C) To assess the reliability of the ancestral sequence reconstructions, the maximum likelihood probability for predicted residues in the ancestral proteins are shown against the relative frequency or fraction of all residues for a given protein.

The multiple sequence alignment and the phylogenetic relationships were then used to infer 171 ancestral sequences for the nodes of the phylogenetic tree. To characterize the changes in NQO1 and NQO2 from a common ancestor to the extant proteins, 5 predicted sequences were chosen for reconstruction (Figure 1A). Three of the sequences included the common ancestor (CA) and the two nodes immediately following the duplication (CA1NQO1 and CA1NQO2). An additional two nodes were chosen at the top of the phylogenetic trees for the amniotes (CA2NQO1 and CA2NQO2). For a given reconstructed ancestral sequence, each amino acyl residue had a particular posterior probability that gave the likelihood of the residue at that position; these results are summarized in Figure 1C for the 5 reconstructed sequences, and the complete data are provided in the Supplemental Tables S1 to S5. The resulting 5 sequences were reliably predicted, with 60% of the residues in the common ancestor (CA) having a probability of 0.95 to 1.0, rising to 90% for the second common ancestor of NQO2 (“CA2NQO2”, Figure 1C). It is notable that the only gap in the alignment of the NQO2 sequences occurs in the C-terminal domain of the common ancestor.

### Purification and Characterization of the Ancestral Proteins

The predicted protein sequences were reverse translated to their corresponding nucleotide sequences for expression in *E. coli*. Protein expression and purification followed our established method for extant NQO2 sequences (Kevin Ka Ki Leung, Litchfield, and Shilton 2012). The ancestral sequences were efficiently expressed in *E. coli* and purified to homogeneity using Ni-NTA affinity chromatography, removal of the affinity tag using TEV protease, and then ion-exchange and gel filtration chromatography. The last step included reconstitution with FAD since our experience indicated that NQO2 produced in *E. coli* contained sub-stoichiometric amounts of FAD (Kevin Ka Ki Leung, Litchfield, and Shilton 2012). The final protein preparations were analyzed for purity by SDS-PAGE (Figure 2A).

**Figure 2.**
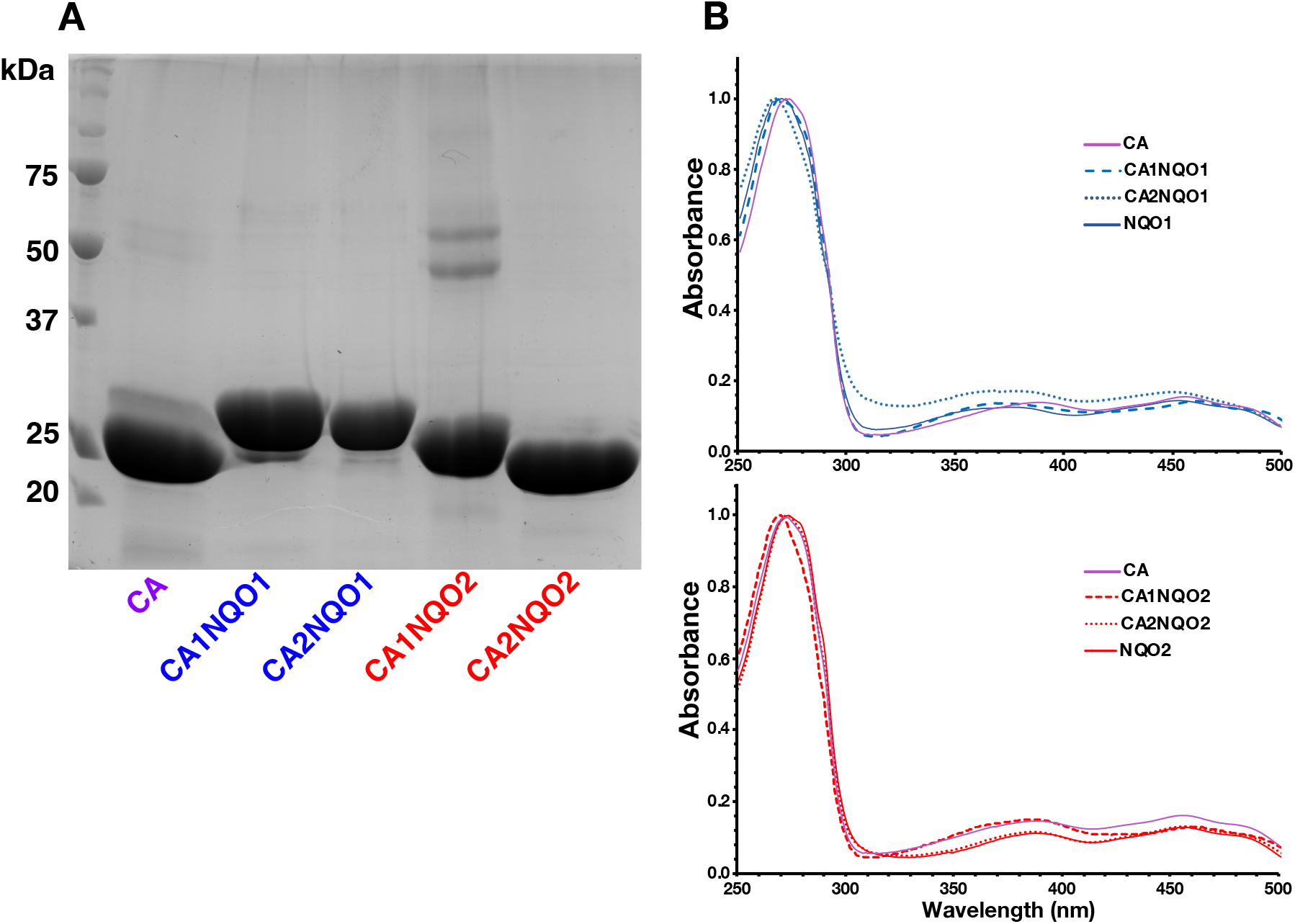
Resurrection and Characterization of Ancestral Quinone Reductases. (A) SDS-PAGE of the purified ancestral proteins. CA is the common ancestor that was duplicated to produce CA1NQO1 and CA1NQO2, the first ancestral NQO proteins in the phylogenetic tree. CA2NQO1 and CA2NQO2 are the ancestral NQO1 and NQO2 proteins at the top of the phylogenetic tree for the amniotes. (B) UV-Visible absorbance spectra of the purified quinone reductases to verify the presence of FAD (maxima at 380 and 455 nm) in the constructs.

The common ancestor (CA) was subject to partial proteolysis during purification (Figure 2A). The predicted C-terminal sequence in the CA contained a two-residue gap and several residues that were predicted with a low probability compared to CA1NQO1 and CA2NQO1 (Figure 1B). Furthermore, predictions of the CA sequence did not always include the C-terminal domain. Since the C-terminal domain is not present in other quinone reductases, it is likely that the CA did not have this domain, and that it was acquired after the gene duplication. On this basis, we suspect that the CA C-terminal domain was disordered and subject to proteolysis. For CA1NQO1, the C-terminal domain was also subject to a small amount of proteolysis (Figure 2A) although posterior probabilities were higher for this sequence and the proteolysis was much more limited that what was observed for the CA.

The preparations of ancestral quinone reductases were further analyzed for FAD content and stability. In a previous study we found that NQO2 overexpressed in *E. coli* was catalytically active but contained only 0.38 equivalents of FMN rather than 1 equivalent of FAD per protomer. Based on this observation we incorporated a reconstitution step in the purification of NQO2 to fully charge the protein with FAD (Kevin Ka Ki Leung, Litchfield, and Shilton 2012). This reconstitution step was incorporated in the purification of all 5 ancestral enzymes, and UV/Vis spectroscopy was used to assess their FAD content. For NQO2 saturated with FAD, the absorbance maximum was at 273 nm (as opposed to 280 nm for NQO2 with 0.38 equivalents of FMN) and the A273/A455 ratio was 7.2 (Kevin Ka Ki Leung, Litchfield, and Shilton 2012). The absorbance spectra of the ancestral proteins indicated maxima between 270 and 275 nm and maxima for the isoalloxazine ring at wavelengths of 455 nm and 380 nm (Figure 2C), with A273/A455 ratios between 6.3 and 7.6, consistent with stoichiometric FAD content.

To assess the stability of the ancestral enzyme constructs, we determined their melting temperature by thermal shift assay, which measures temperature-dependent protein unfolding using a fluorescent dye. The melting temperatures (*T*_M_) were derived from the thermal denaturation profiles using both a fluorescence derivative plot and non-linear fitting to Boltzmann sigmoidal curve, which showed close agreement (Vivoli et al. 2014). The mean *T*_M_ values from fluorescence derivative plots are summarized in Table 1. Generally, the ancestral sequences were more thermostable by at least 10 °C than the corresponding extant proteins. The increased thermostability of the ancestral constructs can be attributed to the use of consensus sequences (Trudeau, Kaltenbach, and Tawfik 2016). Extant NQO2 has a higher thermal stability (*T*_M_ of 64.2°) than extant NQO1 (*T*_M_ of 58.1°) and this greater thermal stability is also present in the NQO2 common ancestors, CA1NQO2 and CA2NQO2, when compared with their ancestral NQO1 orthologues, CA1NQO1 and CA2NQO2 (Table 1).

**Table 1.** Thermal stability of ancestral and extant quinone reductases.

| **Protein** | **$T_m$ (C°)<sup>a</sup>** |
| --- | --- |
| CA | 68.3<br>(67.7 – 68.9) |
| CA1NQO1 | 69.1<br>(64.7 – 73.5) |
| CA2NQO1 | 69.2<br>(68.7 – 69.7) |
| NQO1 | 58.1<br>(56.0 – 60.2) |
| CA1NQO2 | 78.9<br>(78.4 – 79.3) |
| CA2NQO2 | 74.7<br>(74.0 – 75.5) |
| NQO2 | 64.2<br>(62.7 – 65.7) |

<sup>a</sup>Melting temperatures were determined from a thermal shift assay using the fluorescence derivative plot and represent the average of three determinations. Values in parentheses are the upper and lower limits of asymmetric 95% confidence intervals.

### Evolution of the catalytic efficiency of NQO1 and NQO2 with NADH

To trace the molecular evolution of NQO1 and NQO2 cosubstrate specificity, we conducted steady-state kinetic analyses of the ancestral proteins with three different reducing cosubstrates: NADH, NRH, and BNAH. We reported previously that extant NQO2 had catalytic efficiencies with conventional reducing cosubstrates, NAD(P)H, that are almost a million-fold lower than NQO1 (Islam et al. 2022). Notably, NQO2 was incredibly inefficient with NADH, with a catalytic efficiency of 3.0 ± 1.8 M^−1^•s^−1^, comprised of a low *k_cat_* 0.0792 ± 0.018 s^-1^ and a very high *K_M_* of 27 ± 10 mM. In contrast, human NQO1 could use NADH with a catalytic efficiency of 8.3•10^5^ M^−1^ • s^-1^, incorporating a *k_cat_* of 240 ± 12 s^-1^ and *K_M_* of 290 ± 48 µM for NADH in its reaction with menadione.

Through the kinetic analyses we found that NQO1 and NQO2 lineages evolved very different cosubstrate specificities, providing evidence for functional divergence following gene duplication. The common ancestor, CA, had a turnover number for NADH of 5.95 s^-1^ and a modest catalytic efficiency of 9554 M^-1^• s^-1^. Going down the NQO1 lineage, the catalytic efficiency with NADH *increased* first ∼10-fold with CA1NQO1 and then an additional ∼50-fold with CA2NQO1. In contrast, the catalytic efficiency of the NQO2 lineage with NADH *decreased* over 200-fold from the CA to CA1NQO2 and then an additional 15-fold to extant NQO2 (Figure 3A). The inability to use NADH by the NQO2 enzymes was not only due to a decrease in *k_cat_*, but also a large increase in the *K*_M_, into the millimolar range such that it is difficult to measure accurately (Table 2). Extant NQO1 can use both NADH and NADPH almost equally efficiently. We also tested the ability of the ancestral proteins to use NADPH as a reducing cosubstrate to find that the CA and NQO1 ancestors could use NADPH equally efficiently. At the same time, NQO2 enzymes lost the ability to use NADPH. Interestingly, all the ancestral proteins were conserved in their ability to reduce menadione, and the *K*_M_ for menadione was consistent at ∼20 µM throughout the ancestors. However, menadione was found to be a substrate inhibitor for the reducing cosubstrate, and the *K*_I_ for menadione varied between the five ancestors. To summarize, in approximately 450 million years of evolution from a common ancestor, NQO1 acquired a 100-fold increase in catalytic efficiency with NADH, while NQO2 exhibited a 3000-fold decrease in catalytic efficiency with the same co-substrate.

**Figure 3.**
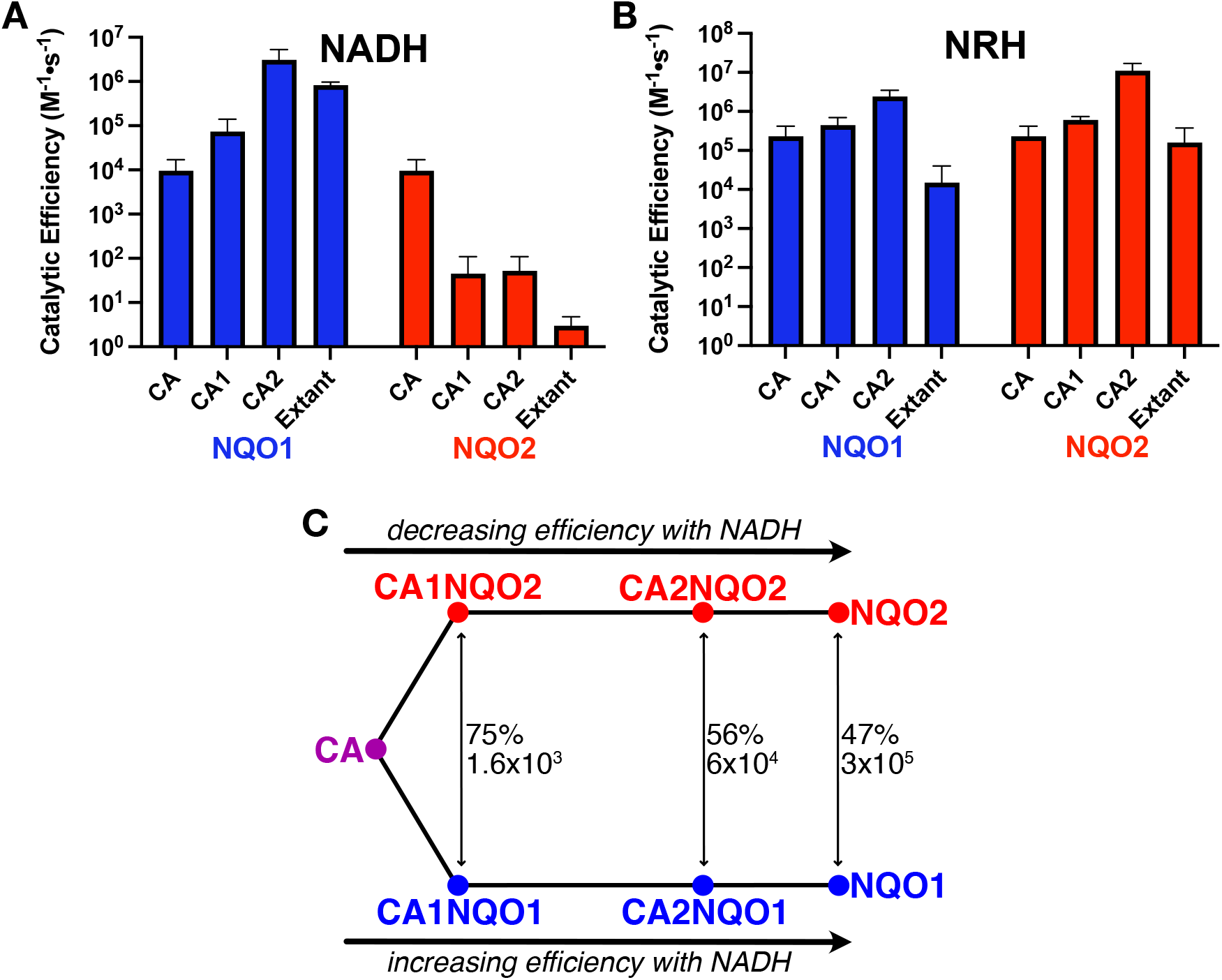
Evolution of Quinone Reductase Cosubstrate Efficiency. The resurrected ancestral and extant quinone reductases were tested for their ability to use the cosubstrates NADH (A) or NRH (B). In both cases, steady-state kinetic analyses included menadione as the quinone substrate. Tables 2, 3, and 4 have the kinetic parameters for NADH, NRH, and BNAH. (C) A graphic illustrating the divergent evolution of the two enzymes from the common ancestor (CA). The numbers indicate the percentage sequence identities and ratio of catalytic efficiencies with NADH of the NQO1 and NQO2 enzymes ([efficiency(NQO1)]/[efficiency(NQO2]) and their two ancestors, CA1 and CA2.

**Table 2.** Kinetic parameters of quinone reductases with NADH.

| Protein | $k_{\text{cat, NADH}}$<br>(s<sup>-1</sup>)<sup>a</sup> | $K_{\text{M, NADH}}$<br>(μM)<sup>a</sup> | $k_{\text{cat}}/K_{\text{M, NADH}}$<br>(M<sup>-1</sup> · s<sup>-1</sup>)<sup>a</sup> |
| --- | --- | --- | --- |
| CA | 5.95<br>(4.50 – 8.64) | 623<br>(414 – 1007) | 9550<br>(5600 – 17000) |
| CA1NQO1 | 59.8<br>(45.4 – 84.4) | 811<br>(493 – 1435) | $7.4 \cdot 10^4$<br>($4 \cdot 10^4$ – $1.4 \cdot 10^5$) |
| CA2NQO1 | 1900<br>(1470 – 2660) | 608<br>(387 – 942) | $3.1 \cdot 10^6$<br>($1.8 \cdot 10^6$ – $5.3 \cdot 10^6$) |
| CA1NQO2 | 0.156<br>(0.109 – 0.286) | 3480<sup>b</sup><br>(2100 – 7840) | 45<br>(22 – 110) |
| CA2NQO2 | 0.299<br>(0.215 – 0.482) | 5790<sup>b</sup><br>(3420 – 11300) | 52<br>(26 – 110) |
| NQO2<sup>c</sup> | $0.0792 \pm 0.018$ | $26800 \pm 10000$<sup>b</sup> | 3.0<br>(1.2 – 4.8) |
| NQO1<sup>c</sup> | $240 \pm 12$ | $290 \pm 48$ | $8.3 \cdot 10^5$<br>($6.5 \cdot 10^5$ – $1.1 \cdot 10^6$) |

<sup>a</sup>Indicated are the upper and lower limits of asymmetric 95% confidence intervals
<sup>b</sup>For these measurements, NQO2 could not be saturated with the co-substrate, and the lack of plateau in the rate at high concentrations of the cosubstrate estimated rate parameters and errors difficult
<sup>c</sup>The kinetic parameters for NQO1 and NQO2 were determined in a previous publication (Islam *et al.* 2022)

**Table 3.** Kinetics of quinone reductases with NRH as cosubstrate.

| Protein | $k_{\text{cat, NRH}}$<br>(s<sup>-1</sup>)<sup>a</sup> | $K_{\text{M, NRH}}$<br>(μM)<sup>a</sup> | $k_{\text{cat}}/K_{\text{M, NRH}}$<br>(M<sup>-1</sup> · s<sup>-1</sup>) |
| --- | --- | --- | --- |
| CA | 194<br>(160 – 249) | 855<br>(593 – 1299) | $2.3 \cdot 10^5$<br>($1.2 \cdot 10^5$ – $4.2 \cdot 10^5$) |
| CA1NQO1 | 130<br>(113 – 154) | 290<br>(219 – 394) | $4.5 \cdot 10^5$<br>($2.9 \cdot 10^5$ – $7.0 \cdot 10^5$) |
| CA2NQO1 | 952<br>(844 – 1091) | 405<br>(313 – 532) | $2.4 \cdot 10^6$<br>($1.6 \cdot 10^6$ – $3.5 \cdot 10^6$) |
| CA1NQO2 | 70.3<br>(66.4 – 74.6) | 115<br>(101 – 132) | $6.1 \cdot 10^5$<br>($5.0 \cdot 10^5$ – $7.4 \cdot 10^5$) |
| CA2NQO2 | 1780<br>(1610 – 1970) | 158<br>(117 – 212) | $1.1 \cdot 10^7$<br>($7.6 \cdot 10^6$ – $1.7 \cdot 10^7$) |
| NQO2 | 101<br>(78.4 – 145) | 629<br>(382 – 1150) | $1.6 \cdot 10^5$<br>($6.8 \cdot 10^4$ – $3.8 \cdot 10^5$) |
| NQO1 | 17.4<br>(13.0 – 27.0) | 1150<br>(675 – 2280) | $1.5 \cdot 10^4$<br>($5.7 \cdot 10^3$ – $4.0 \cdot 10^4$) |

<sup>a</sup> Indicated are the upper and lower limits of asymmetric 95% confidence intervals

### Efficient use of NRH and BNAH by ancestral and extant quinone reductases

Although extant NQO2 cannot use conventional dihydronicotinamide cosubstrates NAD(P)H efficiently, it can catalyze quinone reduction very efficiently given smaller nicotinamide cosubstrates such as NRH, NMEH or BNAH, among others (Knox et al. 2000). To test the hypothesis that NQO2 has evolved to specifically use small nicotinamide cosubstrates such as NRH (S Liao, Dulaney, and Williams-Ashman 1962), we measured the ability of the ancestral NQO1 and NQO2 enzymes to use NRH with menadione as quinone substrate (Figure 3B and Table 3). We found that the common ancestor and the ancestral NQO1 and NQO2 enzymes could use NRH as a reducing cosubstrate with relatively high catalytic efficiencies, with no evidence of systematic change over time. We also measured the oxidation of BNAH by the ancestral enzymes (Table 4) to find whether both enzymes exhibited similar efficiencies with a synthetic dihydronicotinamide cosubstrate. We found that all the ancestral enzymes could use the small synthetic cosubstrate BNAH with high catalytic efficiencies in the range of 10^6^ to 10^7^ M^-1^ • s^-1^. The *K*_M_ for BNAH ranged from 20 µM to 200 µM, and the *k*_CAT_ was from 158 s^-1^ to 1557 s^−1^ with no clear evidence for systematic evolutionary selection in either protein lineage. There is evidence for the existence of NRH under physiological conditions (Yang et al. 2020; Giroud-Gerbetant et al. 2019), but the lack of a systematic evolutionary changes in NRH efficiency observed in both NQO1 and NQO2 lineages, combined with their ability to use a synthetic co-substrate, BNAH, supports the idea that NQO2 did not evolve to use smaller dihydronicotinamide cosubstrates such as NRH. Instead, it appears that both enzymes can use the smaller reducing cosubstrates adventitiously, and NQO2 has evolved to use the conventional reducing cosubstrates inefficiently.

**Table 4.**
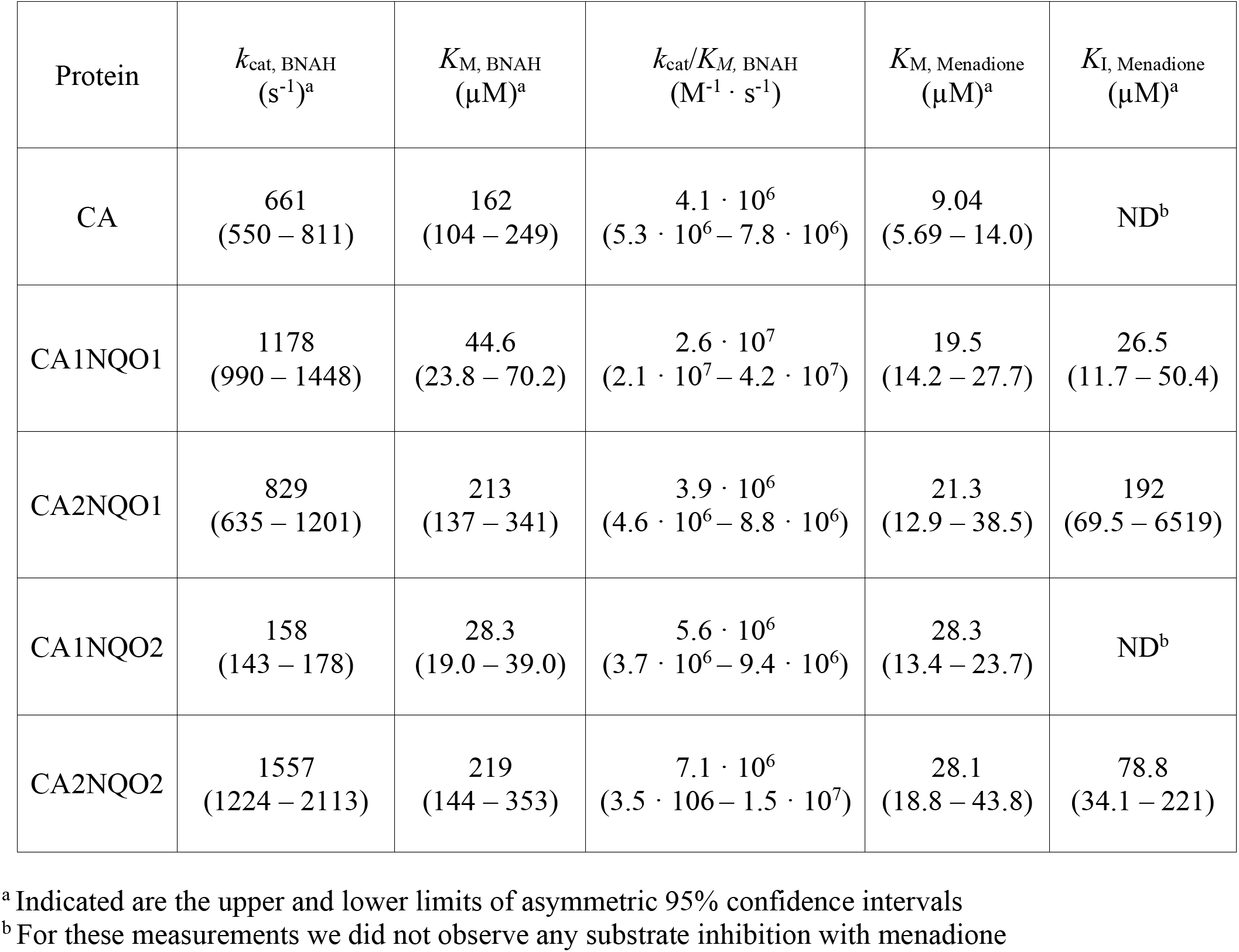
Kinetics of ancestral enzymes with BNAH as cosubstrate.

| Protein | $k_{\text{cat, BNAH}}$<br>(s<sup>-1</sup>)<sup>a</sup> | $K_{\text{M, BNAH}}$<br>(μM)<sup>a</sup> | $k_{\text{cat}}/K_{\text{M, BNAH}}$<br>(M<sup>-1</sup> · s<sup>-1</sup>) | $K_{\text{M, Menadione}}$<br>(μM)<sup>a</sup> | $K_{\text{I, Menadione}}$<br>(μM)<sup>a</sup> |
| --- | --- | --- | --- | --- | --- |
| CA | 661<br>(550 – 811) | 162<br>(104 – 249) | $4.1 \cdot 10^6$<br>($5.3 \cdot 10^6$ – $7.8 \cdot 10^6$) | 9.04<br>(5.69 – 14.0) | ND<sup>b</sup> |
| CA1NQO1 | 1178<br>(990 – 1448) | 44.6<br>(23.8 – 70.2) | $2.6 \cdot 10^7$<br>($2.1 \cdot 10^7$ – $4.2 \cdot 10^7$) | 19.5<br>(14.2 – 27.7) | 26.5<br>(11.7 – 50.4) |
| CA2NQO1 | 829<br>(635 – 1201) | 213<br>(137 – 341) | $3.9 \cdot 10^6$<br>($4.6 \cdot 10^6$ – $8.8 \cdot 10^6$) | 21.3<br>(12.9 – 38.5) | 192<br>(69.5 – 6519) |
| CA1NQO2 | 158<br>(143 – 178) | 28.3<br>(19.0 – 39.0) | $5.6 \cdot 10^6$<br>($3.7 \cdot 10^6$ – $9.4 \cdot 10^6$) | 28.3<br>(13.4 – 23.7) | ND<sup>b</sup> |
| CA2NQO2 | 1557<br>(1224 – 2113) | 219<br>(144 – 353) | $7.1 \cdot 10^6$<br>($3.5 \cdot 10^6$ – $1.5 \cdot 10^7$) | 28.1<br>(18.8 – 43.8) | 78.8<br>(34.1 – 221) |

<sup>a</sup> Indicated are the upper and lower limits of asymmetric 95% confidence intervals
<sup>b</sup> For these measurements we did not observe any substrate inhibition with menadione

**Table 5.**
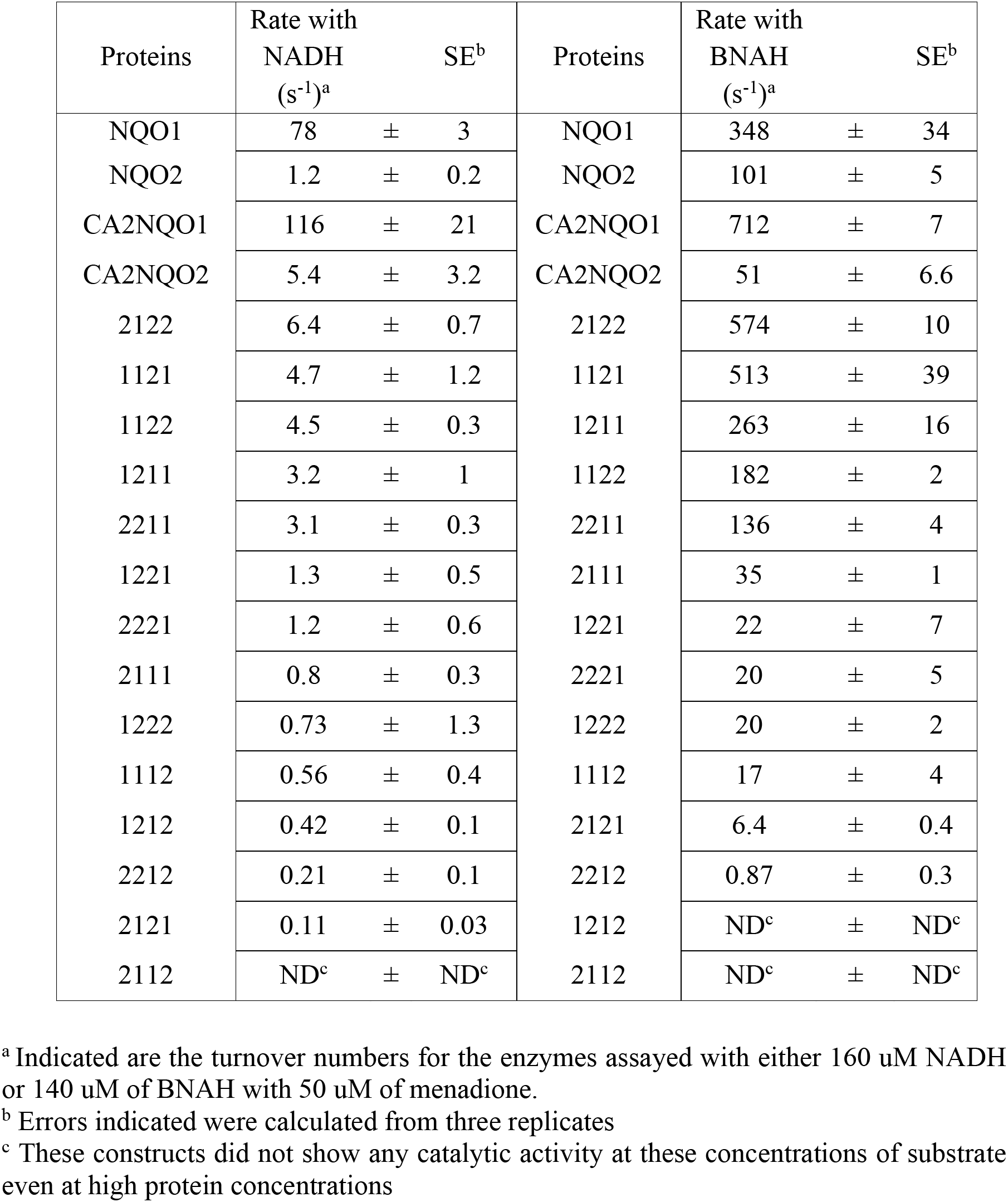
Kinetics of quinone reductase chimeras with NADH and BNAH.

### Molecular determinants of cosubstrate specificity in NQO1 and NQO2

Extant NQO1 and NQO2 differ in their catalytic efficiency with NADH by approximately 300,000-fold, but given their similar structures and 47% sequence identity, it is not clear why NQO2 is not able to use NAD(P)H efficiently. The ancestral sequence reconstruction facilitates an investigation into this issue because the ratio of catalytic efficiencies for the reconstructed enzymes, CA2NQO1:CA2NQO2, is 60000 (Figure 3B), close to the ratio for the extant enzymes, but the reconstructed enzymes have greater homology, with 56% sequence identity representing 103 substitutions in the NQO2 coding region. To assess if there are regions that determine the difference in efficiency with NADH, portions of sequences between the CA2NQO1 and CA2NQO2 were swapped to produce a series of 14 chimeric proteins. The proteins were divided into 4 regions, with breakpoints in conserved sequences, as follows (Figure 4A): segment 1, residues 1 to 100 with 36 substitutions; segment 2, residues 101 to 154 with 19 substitutions; segment 3, residues 155 to 219 with 39 substitutions; and segment 4, residues 220 – 231 of NQO2 with 9 substitutions or the full C-terminal tail (residues 220 – 275) of NQO1. The chimeric quinone reductases were linked to a carbohydrate binding module (CBM) to facilitate single-step, small-scale purification on paper disks (Ball and Basilone 2022). The CBM-NQO[1/2] fusions were expressed in *E. coli* in 2.5 mL cultures, isolated from the cell extracts on paper disks, and NQO[1/2] chimeras were released by incubating the disks with TEV protease. The purified chimeras yielded between 10 and 84 µg of protein; SDS-PAGE analysis of representative preparations is shown in Figure 4B. The chimeras are identified by 4 digits representing the 4 segments of the proteins, with a 1 or 2 for each digit corresponding to the sequence in either CA2NQO1 or CA2NQO2, respectively; for example, the 1121 chimera consists of 3 segments from CA2NQO1 (segment nos. 1, 2, and 4), and one segment (no. 3) from CA2NQO2.

**Figure 4.**
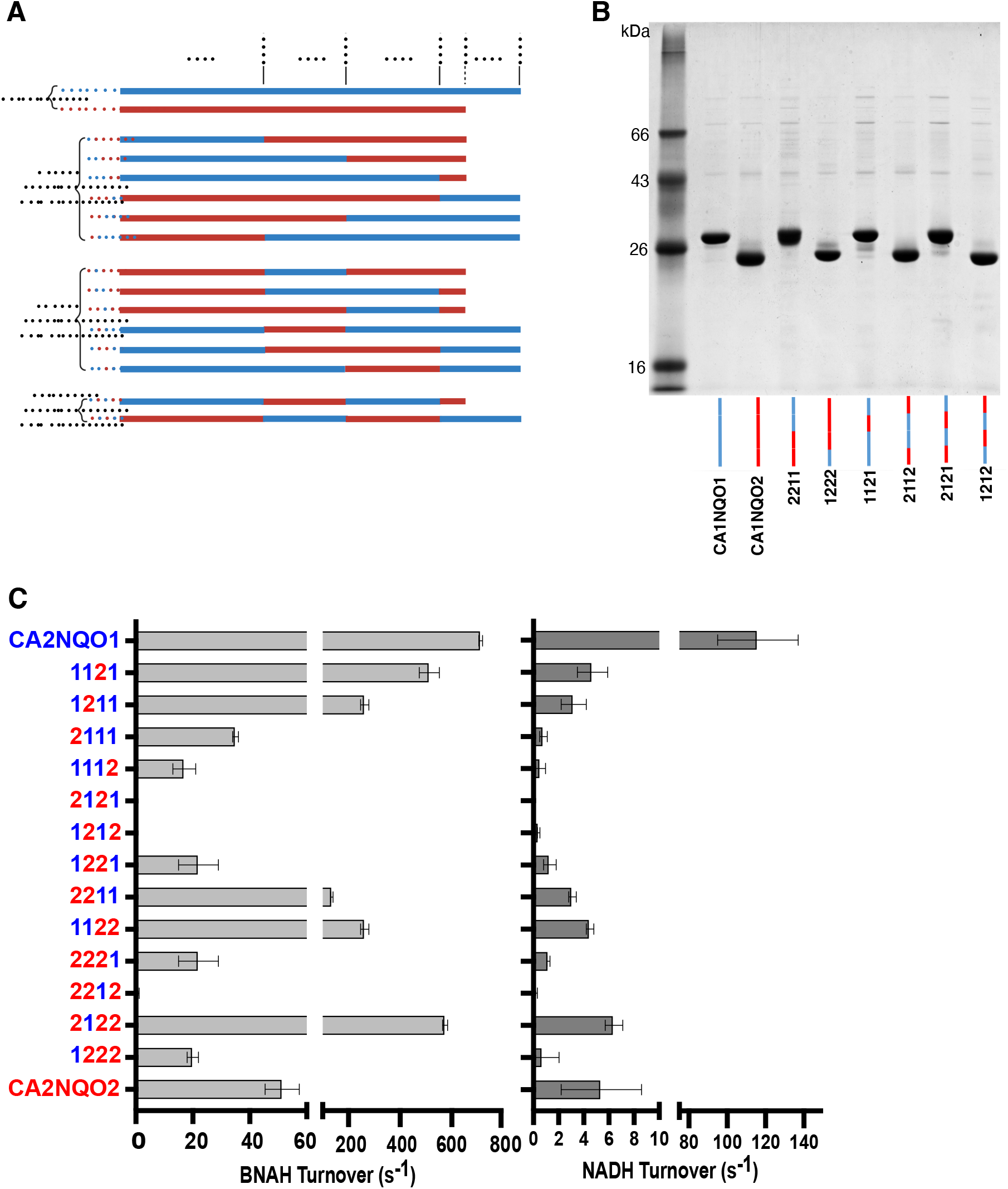
Sequences and purification of chimeric quinone reductase sequences. The sequences of the common ancestors for NQO1 and NQO2, at the top of the phylogenetic tree for the amniotes (i.e. CA2NQO1 and CA2NQO2) were divided into four segments that were then mixed together to produce 14 chimeric proteins. (A) Sequence graphics of the NQO chimeric proteins are provided along with the two parental sequences, CA2NQO1 and CA2NQO2. S1, S2, S3, and S4 are the 4 segments of the parental sequences that were shuffled to produce the chimeras; the breakpoints between segments are indicated by the vertical lines and residue numbers. (B) SDS-PAGE analysis of representative purifications of 8 of the chimeras. The NQO chimeras were fused to a cellulose binding domain (CBD), expressed in small-scale cultures, and purified on paper disks. The NQO chimeras were released from their CBD fusion partner by cleavage with TEV protease. (C) Catalysis by the purified parental and chimeric NQO proteins. The assays included 50 µM menadione and either 140 µM BHAH (left) or 160 µM NADH (right) as cosubstrates, and were normalized for protein concentration to provide the turnover numbers.

Both NQO1 and NQO2 can use BNAH efficiently, and 10 of the 13 purified chimeras exhibited significant catalytic activity with BNAH (Figure 4C), indicating they were properly folded and contained a flavin cofactor. Of note are the constructs that had levels of activity with BNAH that approached that of CA2NQO1, including 1121, 1211, 2211, 1122, and 2122 (Figure 4C), but none of these changes significantly increased the activity with NADH beyond that of CA2NQO2. This is particularly notable for the 1211 and 1121 constructs, which both had robust activity with BNAH but the presence of a single CA2NQO2 segment resulted in almost complete loss of catalytic efficiency with NADH. Overall, one conclusion we can draw from this analysis is that critical residues contributing to the inefficient use of NADH by CA2NQO2 are dispersed throughout the entire sequence.

### Site-directed changes to increase NQO2 efficiency with NADH

To gain further insight into the molecular basis of NQO2 substrate specificity, we used the ancestral sequence analysis to identify residues that could contribute to high catalytic efficiency with NADH. Looking at the NQO1 evolutionary pathway, there was a ∼100-fold increase in catalytic efficiency with NADH in going from the CA to CA2NQO1 (Figure 3A). Sixteen residues were identified that changed going from CA to CA1NQO1 or CA2NQO1, but were conserved in the CA to NQO2 lineage (Figure 5A); these residues could contribute to the *increased* efficiency in NQO1. Therefore, to increase the catalytic efficiency of CA1NQO2, these residues were changed to those observed in CA1NQO1 or CA2NQO1; the changes were P15R, N62D, D128S, F129Y, P130Q, L138F, E154S, P193I, Y195H, E198P, K200A, K202S, N222S, S226N, G227S and Y228N. Conversely, looking at the NQO2 evolutionary pathway, there was a 100-fold decrease in catalytic efficiency going from CA to CA1NQO2 (Figure 3A). Eight residues were identified that were changed going from CA to CA1NQO2 but were conserved in the CA to NQO1 lineage (Figure 5A); these could be residues that contribute to the *decreased* efficiency in NQO2. To increase the catalytic efficiency of CA1NQO2, these residues were changed back to the ones found in the CA; the changes were N24D, S31K, E74L, K166N, Y167V, E194G, C223F, and T224V. In deciding which residues to mutate, we also note that the active site of NQO2 has a negative electrostatic potential compared to the active site of NQO1, which could have important implications for binding the negatively charged phosphates in NAD(P)H. Five of the 26 substitutions introduced positive charges, (P15R, S31K, N162H, Q183K and Y195H) and four neutralized negative charges in NQO2 (E74L, D128S, E154S, and E194G) while two new negative charges were introduced (N24D and N62D). To test the effect of the C-terminal domain of NQO1, a longer mutant NQO2 with the C-terminal domain of CA2NQO1 was also made. The two NQO2CA1 mutant proteins were expressed and purified following the same protocol as for the extant and ancestral proteins; the final preparations were characterized by SDS-PAGE (Figure 5B)

**Figure 5.**
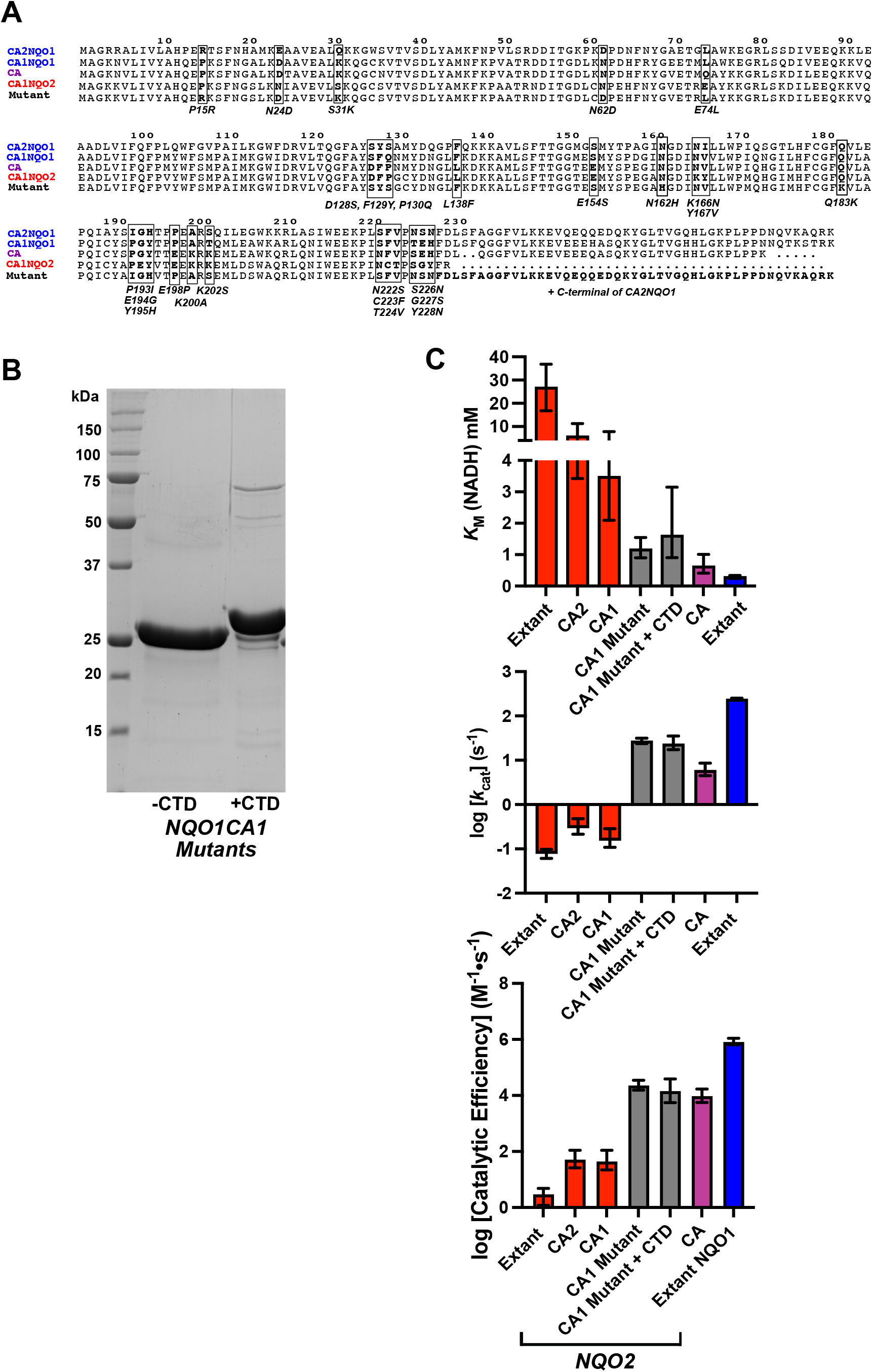
Distributed mutations enhance activity of CA1NQO2 with NADH. (A) The CA1NQO2 ancestor was modified at 26 residues to enhance its activity with NADH; the changes are indicated by boxes and bold type. Most mutations were chosen to replace residues in CA1NQO2 with residues in CA2NQO1 that appeared to be important for NADH catalytic efficiency based on sequence conservation and/or greater positive charge around the active site. Two CA1NQO2 mutants were made, one without a C-terminal domain, and the second with the C-terminal domain of CA2NQO1. (B) SDS-PAGE analysis of the CA1NQO2 mutants. The two CA1NQO2 mutants were expressed and purified following the procedures for histidine-tagged NQO proteins. (C) The kinetic analysis of the mutants indicated a 3-fold decrease in *K*_M_ and 80-fold increase in *k*_CAT_, for an approximate 200-fold increase in catalytic efficiency with NADH.

Our hope was that these mutations, chosen with guidance from our evolutionary analysis, would result in stable proteins that could use NADH more efficiently. In this regard, the CA1NQO2 “short” mutant protein (i.e. no C-terminal domain) was active and exhibited a catalytic efficiency with BNAH (5.6 · 10^7^ M^-1^·s^-1^) that was 10-fold higher than the “parental” CA1NQO2 (5.6 · 10^6^ M^-1^·s^-1^). The CA1NQO2 long mutant (i.e. with the C-terminal domain from CA1NQO1) had a lower catalytic efficiency (2.1 · 10^6^ M^-1^·s^-1^) than the mutant without the C-terminal domain. These results show that the CA1NQO2 mutant proteins were properly folded with bound FAD and retained their catalytic activity with BNAH. The mutations in CA1NQO2 increased its catalytic efficiency with NADH over 200-fold, bringing it back to a level similar to the common ancestor, CA (Figure 5C). The increased catalytic efficiency in the CA1NQO2 mutant proteins (short and long) was primarily due to an 80-fold increase in the *k*_CAT_, along with a 3-fold decrease in *K*_M_ (Figure 5C). Consistent with a previous chimeric construct, where the C-terminal domain of extant NQO1 was added to extant NQO2 (Wu et al. 1997), the presence of the C-terminal domain on the mutant NQO2CA1 enzyme had no significant effect on its ability to use NADH.

### Binding and Efficient use of NADH in NQO1

The significant increase in catalytic efficiency wrought by the changes in the CA1NQO2 mutant prompted us to look more closely at the changes in the protein, and in particular at the electrostatic surfaces, based on the clear difference between NQO1 and NQO2 (Figure 6A). The NQO1 active site has a more positive electrostatic potential around and above FAD cofactor. This region of electrostatic potential above the isoalloxazine ring can be seen in the CA, which uses NADH with moderate efficiency, and in CA1-NQO1, which has an increased efficiency with NADH. However, this region is neutral or slightly negative in CA1-NQO2, which has a drastically decreased catalytic efficiency with NADH as cosubstrate. On the other hand, CA1-NQO2 with the 26 point mutations has a restored basic region, similar to what is observed with the CA, and a similar catalytic efficiency as the CA.

**Figure 6.**
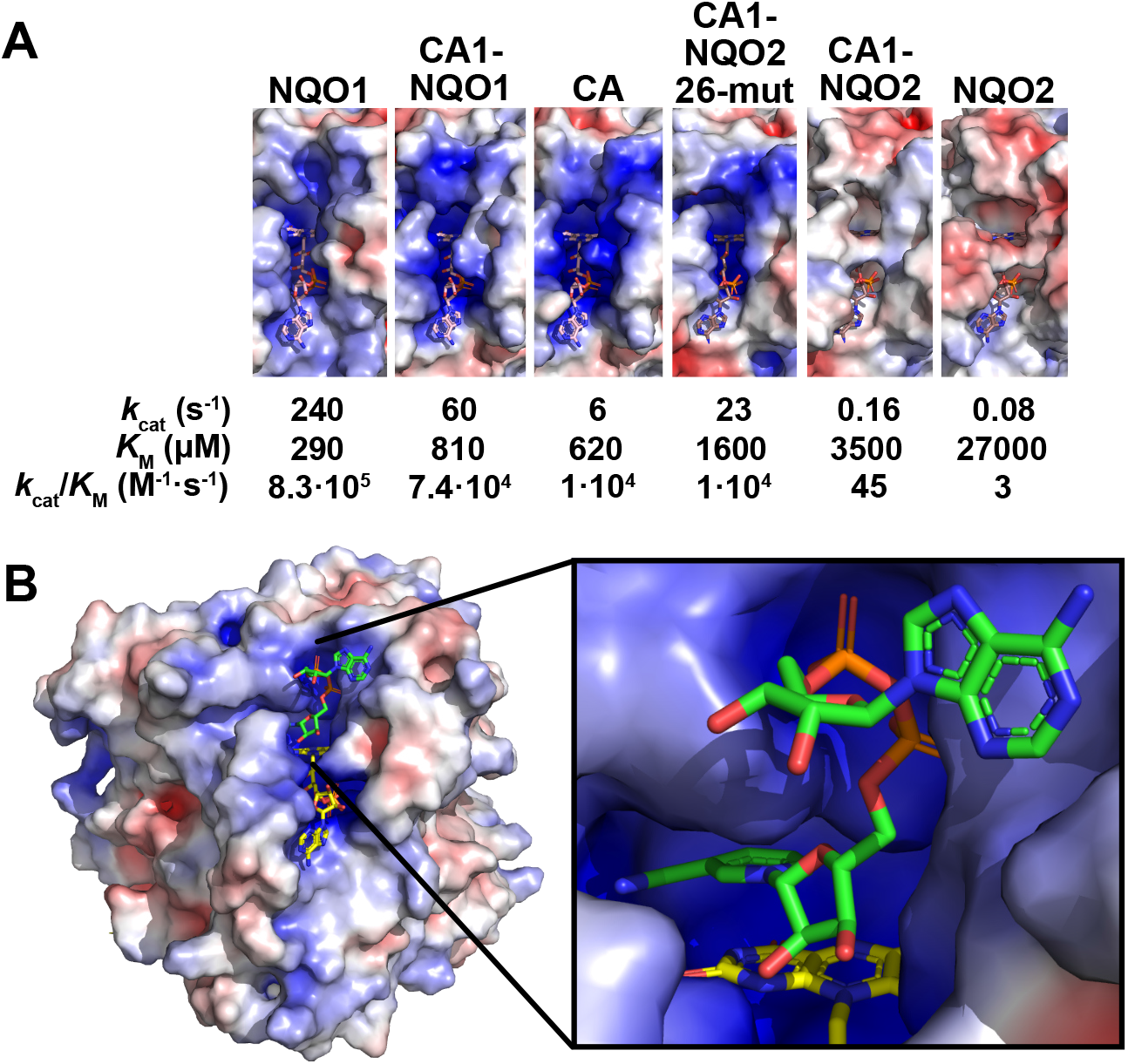
Electrostatics Point to NAD Binding Site in NQO1. (A) The ancestors of NQO1 and NQO2 were homology modelled using crystal structures of the extant enzymes as templates; the CA structure was modelled using NQO2 as a template. The FAD cofactors were included based on their positions in NQO1 and NQO2 structures and provide a common reference point for the surface representations. Electrostatic surfaces for the proteins were calculated using the APBS plugin in PyMol (Baker et al. 2001). The CA uses NADH with moderate catalytic efficiency (*k*_CAT_/*K*_M_ = 10^4^ M^-1^•s^-1^), which is decreased over 200-fold in CA1-NQO2 (*k*_CAT_/*K*_M_ = 45 M^-1^•s^-1^) but regained in the engineered CA1-NQO2 with 26 mutations (CA1-NQO2 26-mut, *k*_CAT_/*K*_M_ = 10^4^ M^-1^•s^-1^). These changes in catalytic efficiency correlate with the disappearance (in CA1-NQO2) and the re-appearance (in CA1-NQO2 26-mut) of the region of positive surface potential above the isoalloxazine ring of FAD. (B) NAD was modelled into the region of positive surface potential of extant NQO1; minor steric clashes were resolved by submitting the complex to a single round of energy minimization using GROMACS (Abraham et al. 2015).

This basic region is clearly present in NQO1, and the correlation between the increase in catalytic efficiency and the appearance of this basic region, in going from CA1NQO2 to the site-directed mutant, suggests that this may represent a binding site for NADH. To investigate this possibility, NAD was modelled into the positive electrostatic region in NQO1, putting the nicotinamide in position for hydride transfer between it and the isoalloxazine ring, fitting the pyrophosphate into a recessed basic pocket, and placing the adenine ring in an indentation on the NQO1 molecular surface. The model complex was subjected to energy minimization using GROMACS (Abraham et al. 2015), fitting very nicely into the site with no steric clashes (Figure 6A).

The positive electrostatic region was present in the CA and on down the NQO1 lineage, but the highly complementary molecular surface for NAD evolved gradually through CA1NQO1 and CA2NQO2. The evolution of this positively charged pocket into which NAD nicely fits correlates with the increasing catalytic efficiency of the enzyme. The modelled conformation of NAD also means that a third phosphate group attached to the 2’-hydroxyl of the adenine ribose is not in contact with the protein, but instead projects into the solvent, consistent with the fact that NQO1 can use both NADH and NADPH equally well (Ernster, Danielson, and Ljunggren 1962).

## Discussion

When Williams-Ashman and co-workers originally purified a redox enzyme, now known as NQO2, that did not catalyze the oxidation of NAD(P)H, but would catalyze the oxidation of NRH or N-propyl-dihydronicotinamide, they speculated that NRH is formed in cells, and the function of the NQO2 enzyme was to oxidize the NRH to NR, thereby avoiding an accumulation of NRH (S Liao, Dulaney, and Williams-Ashman 1962). The ancestral sequence reconstruction shows that the common ancestor of NQO1 and NQO2 had relatively modest catalytic efficiency with NADH and greater efficiency with NRH and the synthetic cosubstrate, BNAH. After the gene duplication, the ancestors of NQO1 and NQO2 retained their ability to use small natural or synthetic dihydronicotinamide cosubstrates but diverged in their use of NAD(P)H, with NQO1 gaining catalytic efficiency, and NQO2 losing catalytic efficiency. We conclude that NQO2 did not evolve to use NRH or other small nicotinamide cosubstrates efficiently, but for an unknown reason has evolved to use NAD(P)H inefficiently. The evolution to use the common cellular dihydronicotinamide cosubstrates inefficiently is unexpected for an enzyme such as NQO2, and this observation may provide clues to cellular functions of NQO2 beyond the efficient catalytic reduction of quinones and other electrophiles.

We are confident that the ancestral sequence reconstruction provided a faithful representation of the evolutionary divergence in cosubstrate specificity. A feature of the reconstructed enzymes that may have an impact on their functional characteristics is that they were about 10°C more thermostable than the extant enzymes. The increased thermal stability is a good indication that the proteins are correctly folded; however, the increased stability and potential changes in protein dynamics could affect catalytic activities. Since we used the same methods to predict the ancestral sequences and obtained the same 10°C shift in both the NQO1 and NQO2 ancestors, it is unlikely that the changes in stability are selectively affecting cosubstrate specificity in one lineage but not the other. Additionally, ancestral proteins reconstructed by maximum likelihood (ML) tend to have higher thermostabilities, not necessarily because the ancestral proteins were more stable, but because of the “consensus effect” and the nature of statistics used for the reconstruction (Trudeau, Kaltenbach, and Tawfik 2016; Furukawa et al. 2020). For example, the tendency of ancestral reconstructed proteins to exhibit higher thermostability is more pronounced for ML over Bayesian Inference (BI) reconstruction. This is because ML methods have higher selectivity for the most common, or “consensus”, amino acids (Williams et al. 2006). ML continues to be used for ancestral state reconstruction and was found to be the most effective method for predicting ancestral sequences correctly on a site-by-site basis (Williams et al. 2006).

The ancestral sequence reconstruction has also pointed towards a binding site for the nicotinamide cofactors in this family of enzymes. The site-directed mutagenesis that successfully increased the catalytic efficiency of CA1-NQO2 highlighted evolutionary changes in the surface electrostatic potential of the NQO1 and NQO2 lineages. These electrostatic changes correlated with changes in catalytic efficiency with NAD(P)H and facilitated modelling of NAD in a putative binding site on NQO1. Although somewhat speculative, the “hand-in-glove” fit of NAD into this site of NQO1, combined with the projection of the adenine ribose into the solvent, which is consistent with the hallmark ability of NQO1 to use both NADH and NADPH as reducing co-substrates, strongly suggests that this may be the bona-fide binding site for NAD(P)H in this family of enzymes. This idea is further supported by the observation that, in addition to a neutral to negative electrostatic potential in the NQO2 lineage, the site is sterically occluded and unable to accommodate the pyrophosphate moiety in NAD(P)H in extant NQO2 and its ancestors, consistent with their very poor catalytic efficiency with NAD(P)H.

One aspect of the ancestral reconstruction subject to uncertainty is the existence of an additional C-terminal domain on the common ancestor (CA). The ∼50 residue NQO1 C-terminal domain is conserved in extant ray-finned fish and mammals, and on this basis the gene duplication must have happened during the evolution of ancient bony fish, prior to their separation into the Actinopterygii (ray-finned fish) and Sarcopterygii (lobe-finned fish, which gave rise to tetrapods). The two possible scenarios for this evolution are that the C-terminal domain was present on the CA and then lost from the NQO2 lineage, or that it was absent from the CA, and then evolved during the early evolution of the NQO1 lineage. Depending on the data and method used for the phylogeny and reconstruction, the NQO1 C-terminal domain could be predicted to be present or not on the CA. We used a reconstruction in which a truncated C-terminal domain was predicted to be present on the CA, however the shorter C-terminal domain on the CA appeared to be highly labile to proteolytic degradation, suggesting an unstable structure. We also noted a tendency of the common ancestor of the NQO1 lineage (i.e. CA1NQO1) to undergo some proteolysis during purification, but not for CA2NQO1 and extant NQO1, consistent with an evolution of this domain down the NQO1 lineage. In addition, the C-terminal domain on NQO1 is unique for the NQO1 enzymes in mammals and extant fish; the bacterial enzymes do not contain a similar C-terminal domain, and in fact are slightly shorter (by about 15 residues) than even NQO2, but they are still able to catalyze the NAD(P)H-dependent reduction of electrophiles. In summary, it appears likely that the C-terminal domain of NQO1 evolved after the initial gene duplication to provide additional physiological function.

NQO1 has documented enzymatic functions in the cell, in particular the transfer of electrons to quinones and other electrophiles (Ross and Siegel 2021). The inefficient use of the common dihydronicotinamide cosubstrates NAD(P)H by NQO2 makes it unlikely that NQO2 has a similar catalytic function as NQO1. In fact, in all cell types tested NQO2 cannot function effectively as a catalyst for reduction of electrophiles unless an exogenous small dihydronicotinamide cosubstrate is supplied to the cell media (Knox et al. 2000; Islam et al. 2022). On this basis, after the gene duplication NQO1 retained the enzymatic function of the ancestral NQO enzyme, but NQO2 underwent neofunctionalization, or the acquisition of a completely different function. Evidence for neofunctionalization of the NQO2 paralog comes from the very different effect of knocking out NQO1 or NQO2 in mice. NQO1 knockout mice demonstrated increased sensitivity to menadione toxicity, consistent with a role for NQO1 in reduction and detoxification of menadione (Radjendirane et al. 1998). In contrast, NQO2 knockout mice showed decreased sensitivity to menadione toxicity, a difference that was even more pronounced when NRH was included in the menadione treatment (D. J. Long et al. 2002). In other words, the presence of NQO2 exacerbates the toxicity of menadione, an effect that is even stronger when a suitable exogenous dihydronicotinamide cosubstrate is supplied. Consistent with these knockout experiments, overexpression of NQO2 in Chinese hamster ovary (CHO) cells increased the toxicity of menadione compared to wild-type cells, and the menadione toxicity was further increased when it was administered in the presence of NRH (Celli et al. 2006). Based on the effect of added NRH, these researchers attributed the adverse effect of NQO2 to metabolic activation of menadione; however, it is not obvious why 2-electron reduction by NQO1 and NQO2 would produce such different cellular outcomes from menadione exposure, with NQO1 decreasing menadione toxicity and NQO2 increasing it. The herbicide paraquat (PQ) creates oxidative stress in mammalian cells that can lead to cell death, and inhibition or knock-down of NQO2 decreases the toxicity of PQ. In contrast to menadione, PQ is neither a substrate nor inhibitor of NQO2; however shRNA silencing or inhibition of NQO2 attenuated cell death induced by paraquat (Janda et al. 2013).

These observations that knockout, silencing, or inhibition of NQO2 increases cell viability under oxidative stress is interesting because NQO2 is frequently discovered in off-target interactions with therapeutic kinase inhibitors. NQO2 was found interacting with bisindolmaleimide protein kinase C inhibitors (Brehmer et al. 2004); the Abl kinase inhibitor imatinib (Bantscheff et al. 2007; Rix et al. 2007); and tetrabromo inhibitors of CK2 (Duncan et al. 2008). In a more comprehensive and systematic search for off-target interactions of kinase inhibitors, NQO2 was found to be interacting with 35 of approximately 240 kinase inhibitors tested (Klaeger et al. 2017). It is noteworthy that NQO1 was not identified as interactor with any of the kinase inhibitors tested, despite its high degree of structural similarity to NQO2. Structural analysis of selected NQO2 and kinase inhibitor complexes indicate binding to the active site of the enzyme, directly on top of the isoalloxazine ring of FAD (Winger et al. 2009; Kevin K. K. Leung and Shilton 2015; Klaeger et al. 2017), a site that bears little if any resemblance to the ATP binding site of kinases, to which almost all the inhibitors are targeted. These observations suggest that NQO2 may somehow contribute to the effects of the kinase inhibitors that are successfully ferried through drug selection processes and clinical trials. For example, given that silencing or inhibition of NQO2 increases cell viability under stress conditions, it may be that the toxicity of certain kinase inhibitors for untransformed cells is decreased if NQO2 is also inhibited. Alternatively, the combined inhibition of a particular kinase pathway and binding to NQO2 may contribute synergistically to toxicity for transformed cells. The mechanism by which binding of a kinase inhibitor to NQO2 could affect its function is unknown, but there are examples of flavoproteins that have a redox signalling function (Becker, Zhu, and Moxley 2011). In this regard, it is also worth considering that small molecules like the kinase inhibitors (Kevin K. K. Leung and Shilton 2015) or the antimalarials primaquine and chloroquine (Kwiek, Haystead, and Rudolph 2004; Kevin Ka Ki Leung 2015) can bind preferentially to either the oxidized or reduced form of the proteins, with implications for a potential signalling function.

In addition to its evolution towards inefficient use of the canonical redox cosubstrates, NAD(P)H, another major difference between NQO1 and NQO2 is the presence of a metal binding site in NQO2 (Foster et al. 1999). This metal site appears to be a type-I copper site, and NQO2 isolated from mammalian tissue contained sub-stoichiometric amounts of copper and zinc (Kwiek, Haystead, and Rudolph 2004). The presence of a conserved copper binding site in NQO2 is interesting in view of observations that NQO2 is a source of reactive oxygen species (ROS) in cells (Cassagnes et al. 2015; Gould, Elkobi, et al. 2020). The presence of NQO2 and its production and/or regulation of cellular ROS have been linked to learning and long-term memory acquisition (Rappaport et al. 2015; Gould, Elkobi, et al. 2020; Gould, Sharma, et al. 2020; Gould et al. 2021), although the involvement of the metal site in these cases has not been explored. Furthermore, NQO1, without a metal site, can also be a source of cellular ROS (Cassagnes et al. 2015). The function of the metal site in NQO2, and whether its presence is somehow linked to its inefficient use of NAD(P)H cosubstrates remains to be determined.

## Supporting information

Supplemental Full Phylogenetic Tree

Tables S1-S5 Ancestral Sequence Probabilities

