## Supplementary figures and images for "Evolutionary analysis of Quinone Reductases 1 and 2 suggest that NQO2 evolved to function as a pseudoenzyme"

### Supplemental Full Phylogenetic Tree

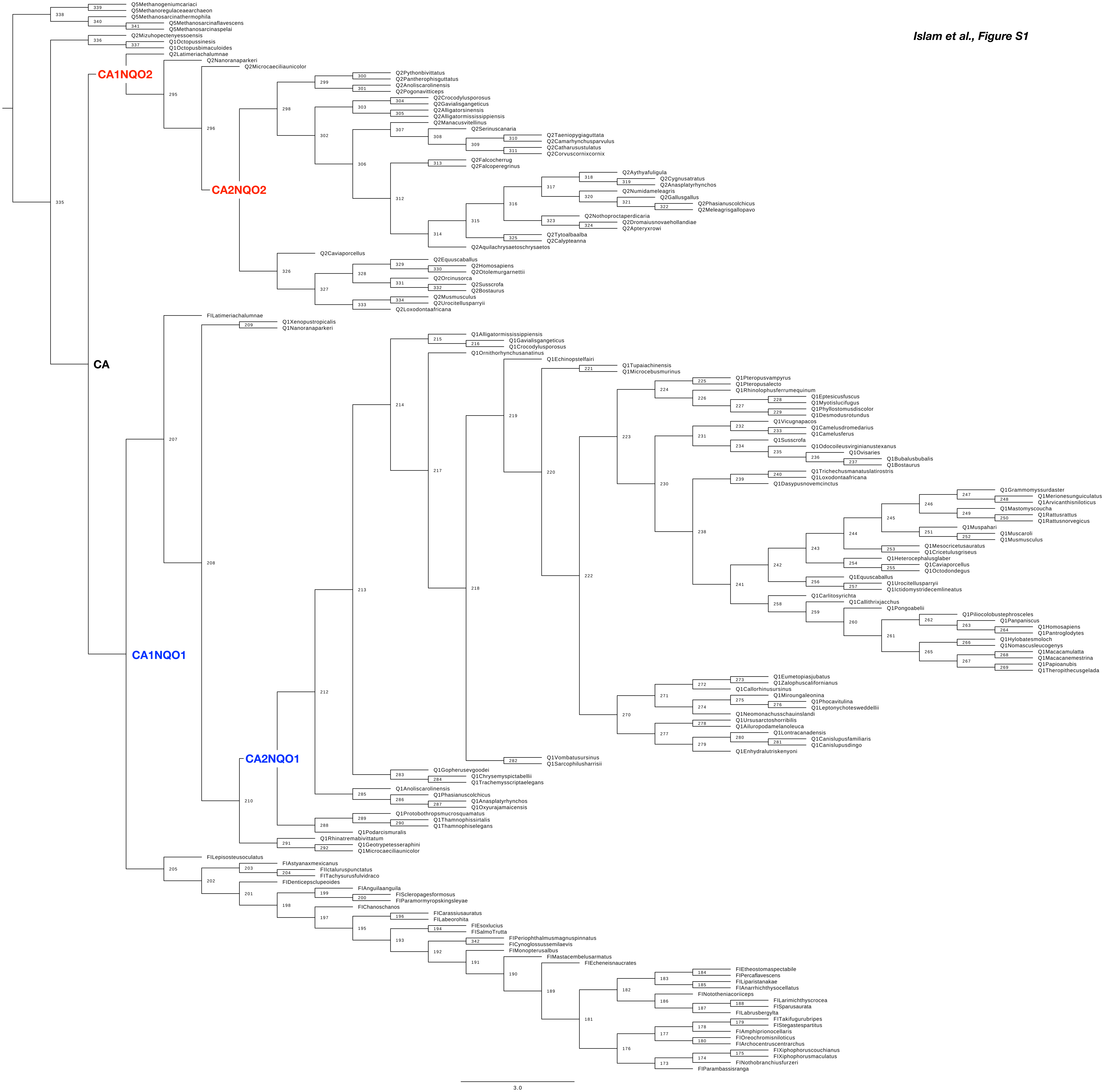


*Islam et al., Figure S1*
